## Supplementary material for "MGSurvE: A framework to optimize trap placement for genetic surveillance of mosquito population": Mathematical description of the model

### MGSurvE: A framework to optimize trap placement for genetic surveillance of mosquito population
### S1 Text: Model Description

Héctor M. Sánchez C.[1*], David L. Smith [2,3], John M. Marshall [1]

**1** Divisions of Epidemiology and Biostatistics, School of Public Health, University of California, Berkeley, California, United States of America
**2** Institute for Health Metrics and Evaluation, University of Washington, Seattle, WA, USA
**3** Department of Health Metrics Sciences, School of Medicine, University of Washington, Seattle, WA, USA

*

#### Mathematical Formulation

The default implementation of MGSurvE assumes individuals traverse a landscape following "random-walker" behavior according to a movement matrix, $\tau$, the entries of which, $\tau_{ij}$, describe the probability of moving from population $i$ to population $j$ in one time step. Incorporating traps as absorbing states within this formulation allows us to calculate outcomes of interest, such as the mean time to absorption (i.e., being trapped) using the properties of absorbing Markov chains.

##### Movement

The movement weight, $\alpha$, between two sites, $s_i \to s_j$, is calculated according to an arbitrary shape parameter, $\kappa$, that is a function of the distance, $d$, between them, and biological modifying parameters, $\rho$. To account for resource-directed movement, this shape parameter is then multiplied by the probability of movement, $\lambda$, from the source site-type, $\hat{s}_i$, to the destination's site-type, $\hat{s}_j$:

$$\alpha(s_i \to s_j) = \kappa(d(s_i \to s_j), \rho) * \lambda(\hat{s}_i, \hat{s}_j) \tag{1}$$

We calculate the movement weights between all the sites in the landscape and normalize the resulting matrix to obtain our migration matrix $\tau$:

$$\tau_{s_n \times s_n} = \begin{pmatrix} \alpha(s_1 \to s_1) & \alpha(s_1 \to s_2) & ... & \alpha(s_1 \to s_{n-1}) & \alpha(s_1 \to s_n) \\ \alpha(s_2 \to s_1) & \alpha(s_2 \to s_2) & ... & \alpha(s_2 \to s_{n-1}) & \alpha(s_2 \to s_n) \\ \vdots & & \ddots & & \vdots \\ \alpha(s_{n-1} \to s_1) & \alpha(s_{n-1} \to s_2) & ... & \alpha(s_{n-1} \to s_{n-1}) & \alpha(s_{n-1} \to s_n) \\ \alpha(s_n \to s_1) & \alpha(s_n \to s_2) & ... & \alpha(s_n \to s_{n-1}) & \alpha(s_n \to s_n) \end{pmatrix} \tag{2}$$

All rows of $\tau$ sum to 1, as individuals are limited to move within the $s_n$ set of sites.

#### Movement with Traps

For the movement matrix incorporating traps as absorbing states, $\chi$, we follow a similar process to calculate the trap probability weights, $\delta$, as we did for the movement weights, $\alpha$. We determine the movement probability from any given site to any given trap based on the attractiveness profile, $\hat{\eta}$, and the trap-type attractiveness relative to the current mosquito resource, $\phi(\hat{s_i}, \hat{t_j})$. Equations 4 and 5 are included to mathematically describe that traps are absorbing states:

$$\delta(s_i \rightarrow t_j) = \hat{\eta}(d(s_i \rightarrow t_j), \hat{\rho}_{trap}) * \phi(\hat{s_i}, \hat{t_j}) \quad \Rightarrow \nu_{s_n \times t_n} \tag{3}$$

$$\delta(t_i \rightarrow s_j || t_i \rightarrow t_j | i \neq j) = 0 \quad \Rightarrow 0_{t_n \times s_n} \tag{4}$$

$$\delta(t_i \rightarrow t_i) = 1 \quad \Rightarrow I_{t_n \times t_n} \tag{5}$$

With these pieces in place, we can define our normalized full movement block matrix with traps as follows, noting that rows must be renormalized to account for possible movement from a site to a trap or to another site:

$$\chi_{(s_n+t_n)\times(s_n+t_n)} = \begin{pmatrix} \tau_{s_n \times s_n} & \nu_{s_n \times t_n} \\ 0_{t_n \times s_n} & I_{t_n \times t_n} \end{pmatrix} \tag{6}$$

#### Fitness Function

With the Markov chain transition matrix including absorbing traps in canonical form, $\chi$, we can use the fundamental matrix, $F$, to calculate a range of Markov chaing properties, including the expected number of time steps before an individual is trapped/absorbed. The fundamental matrix, $F$, is given by:

$$F_{s_n \times s_n} = (I - \tau_{s_n \times s_n})^{-1} \tag{7}$$

The $F$ matrix contains information on how many time steps an individual is expected to spend in site $j$ given that it started in site $i$, before falling into a trap.

Finally, to compute the fitness value $\phi$, we calculate the expected number of time steps before being trapped, when starting in transient site $i$:

$$t_{s_n \times 1} = F_{s_n \times s_n} * 1_{s_n \times 1} \tag{8}$$

Two metrics that we use in the current paper are: i) the expected number of time steps before being trapped, averaged over all origin sites, i.e., $mean(t_{s_n \times 1})$, and ii) the maximum expected number of time steps before being trapped, considering all origin sites, i.e., $max(t_{s_n \times 1})$. The latter case represents the worst case scenario.
