## Supplementary material for "MGSurvE: A framework to optimize trap placement for genetic surveillance of mosquito population": Demonstrations code

MGSurvE: A framework to optimize trap placement for genetic surveillance of mosquito population
S2 Text: Urban and Semi-Rural Landscape Code

Héctor M. Sánchez C.[1*], David L. Smith [2,3], John M. Marshall [1]

**1** Divisions of Epidemiology and Biostatistics, School of Public Health, University of California, Berkeley, California, United States of America
**2** Institute for Health Metrics and Evaluation, University of Washington, Seattle, WA, USA
**3** Department of Health Metrics Sciences, School of Medicine, University of Washington, Seattle, WA, USA

*

In this document we provide a summary of the needed code to run the examples shown in the manuscript. For more information, and tutorials have a look at our documentation: https://chipdelmal.github.io/MGSurvE; and to the source code of these demonstrations:
https://github.com/Chipdelmal/MGSurvE/tree/main/MGSurvE/demos.

### Urban Landscape: "Yorkey's Knob"and "Trinity Park"

To optimize the location of our traps, the first thing we need to do is to load our pointset, which is stored in *lon/lat* coordinates in a CSV file. We can import it as follows:

```
# Load pointset and set all sites to the same point-type (0)
YK_LL = pd.read_csv(LND_PTH, names=['lon', 'lat'])
YK_LL['t'] = [0]*YK_LL.shape[0]
# Define movement kernel to an Aedes-aegypti-like species
mKer = {
    'kernelFunction': srv.zeroInflatedExponentialKernel,
    'kernelParams': {'params': srv.AEDES_EXP_PARAMS, 'zeroInflation': 1-0.28}
}
```

With this in place, we can setup the attractiveness kernels of the traps, and define how many we will want of each type. The attractiveness kernels are defined in a dictionary where each entry contains a kernel (a function that maps distance and some constants into a probability), and its shape parameters:

```
# Setting up trap attractiveness kernels
tKer = {
    1: {
        'kernel': srv.sigmoidDecay,
        'params': {'A': 0.5, 'rate': .25, 'x0': 1/0.0629534}
    },
    0: {
        'kernel': srv.exponentialDecay,
        'params': {'A': 0.5, 'b': 0.0629534}
```

```
    }
}
# Define of traps and their type (as defined by the kernel dictionary)
nullTraps = [0]*TRPS_NUM
cntr = (
    [np.mean(YK_LL['lon'])]*TRPS_NUM,
    [np.mean(YK_LL['lat'])]*TRPS_NUM
)
sid = [0]*TRPS_NUM
traps = pd.DataFrame({
    'sid': sid,
    'lon': cntr[0], 'lat': cntr[1],
    't': TRAP_TYP, 'f': nullTraps
})
```

With the traps in place, we can setup our landscape object. This is done by passing all the information of the previous steps to its initialization routine (for more information look at the object's documentation entry https://chipdelmal.github.io/MGSurvE/build/html/MGSurvE.html#MGSurvE.landscape.Landscape):

```
# Setup landscape up with point-sites and traps
lnd = srv.Landscape(
    YK_LL,
    kernelFunction=mKer['kernelFunction'],
    kernelParams=mKer['kernelParams'],
    traps=traps, trapsKernels=tKer
)
```

And finally, we can optimize it with our simplified wrapper function:

```
# Optimize the landscape with automatic parameters
(lnd, logbook) = srv.optimizeTrapsGA(
    lnd, generations=GENS,
    pop_size='auto', selection_params='auto',
    mating_params='auto', mutation_params='auto',
    fitFuns={'inner': np.sum, 'outer': np.mean}
)
# Export results to disk
srv.exportLog(logbook, OUT_PTH, 'YKN_LOG')
srv.dumpLandscape(lnd, OUT_PTH, 'YKN_LND', fExt='pkl')
```

Which runs the algorithm for the defined number of generations and exports the positions into a text file. The full running code can be downloaded from: https://github.com/Chipdelmal/MGSurvE/blob/main/MGSurvE/demos/YKN/YKN-Discrete.py; and is also included with the package installation.

### Semi-Rural Landscape: "São Tomé"

In this application we will set to two fixed traps and import an externally-generated migration matrix. To get started, we load our point-set the same way we did before, along with our migration file:

```
# Load pointset
SAO_TOME_LL = pd.read_csv(path.join('./GEO', 'STP_LatLonN.csv'))
# Load migration matrix
migration = np.genfromtxt(
    path.join('./GEO', 'STP_MigrationN.csv'),
    delimiter=','
)
SAO_TOME_MIG = normalize(migration, axis=1, norm='l1')
# Bounding box and ids of the sites where we want fixed traps
SAO_LIMITS = ((6.41, 6.79), (-0.0475, .45))
SAO_FIXED = [24, 212]
```

Now, to do our optimization, we need to pass a *sid* parameter to our traps dataframe. This number represents the identifier of the site at which each trap will be located (all but two traps will be initialized at the site 0 as they'll be randomized in the optimization cycle, whereas the other two will be initialized at the ids 24 and 212):

```
sid = [0]*(TRPS_NUM-FXD_NUM) + SAO_FIXED
nullPos = [0]*TRPS_NUM
traps = pd.DataFrame({
    'sid': sid,
    'lon': nullPos, 'lat': nullPos,
    't': initTyp, 'f': initFxd
})
tKer = {0: {'kernel': srv.exponentialDecay, 'params': {'A': 1, 'b': .0075}}}
```

We can now initialize our landscape:

```
lnd = srv.Landscape(
    SAO_TOME_LL,
    migrationMatrix=SAO_TOME_MIG,
    traps=traps, trapsKernels=tKer,
    landLimits=SAO_LIMITS
)
lndGA = deepcopy(lnd)
```

Up to this point, we have followed almost the same procedure as we did for the previous example, with the slight changes necessary to defined immovable traps, discrete traps locations, and external migration kernels. Now we call our wrapper optimization function:

```
(lnd, logbook) = srv.optimizeDiscreteTrapsGA(
    lndGA, generations=GENS,
    pop_size='auto', selection_params='auto',
    mating_params='auto', mutation_params='auto',
    fitFuns={'inner': np.sum, 'outer': np.mean}
)
```

which returns our landscape with the optimized traps in place. The full running code is available at https://github.com/Chipdelmal/MGSurvE/blob/main/MGSurvE/demos/STP/STP-Discrete.py.
